# Evaluation of US state pollinator plans using 3 evidence-based policymaking frameworks

**DOI:** 10.1101/2021.06.15.447774

**Authors:** Kaitlin Stack Whitney, Briana Burt Stringer

## Abstract

After the US federal government created a national pollinator protection plan in 2015, many states followed with their own. Since their goal is to promote pollinating insect conservation, we wanted to know whether the state plans are using best practices for evidence-based science policy. In early 2019 we found and downloaded every existing, publicly available US state pollinator protection plan. We then used content analysis to assess the goals, scope, and implementation of state-level pollinator protection plans across the US. This analysis was conducted using three distinct frameworks for evidence-based policymaking: US Department of Interior Adaptive Resources Management (ARM), US Environmental Protection Agency management pollinator protection plan (MP3) guidance, and Pew Trusts Pew-MacAthur Results First Project elements of evidence-based state policymaking (PEW) framework. Then we scored them using the framework criteria, to assess whether the plans were using known best practices for evidence based policymaking. Of the 31 states with a state pollinator plan, Connecticut was the state with the lowest total score across the three evaluation frameworks. The state with the highest overall scores, across the three frameworks, was Missouri. Most states did not score highly on the majority of the frameworks. Overall, many state plans were lacking policy elements that address monitoring, evaluation, and adjustment. These missing elements impact states’ ability to achieve their conservation goals. Our results indicate that states can improve their pollinator conservation policies to better match evidence-based science policy guidance, regardless of which framework is used.

## Introduction

Pollinators are critical to healthy, functioning ecosystems. Pollinators are animals that carry pollen from male to female flowers, on the same or different flower or plant. This interaction between flowering plants and animals exists across many generalist and specialist taxa, and it is estimated to have existed for over 100 million years (van der Koii and Ollerton 2020). An estimated 87.5% of wildflowers in temperate and tropical regions depend on pollinating animals (Ollerton et al. 2011), so pollinators are critical to the maintenance of pollination, a regulating ecosystem service.

Many pollinators are insects. Honey bees (*Apis mellifera*) are one of the most studied and celebrated insect pollinators in human history. They are valued for their harvestable honey and beeswax products, as well as their pollination of crop and wild plants (Hung et al. 2018). Yet honey bees are only one species of bee. Wild bees abound - in New York alone, there are an estimated 416 species of bee (Cornell n.d). Honey bees are also managed pollinators, which means they are raised as domestic animals, like livestock. Honey bee colonies are bred and maintained by human keepers. Wild bees, in contrast, are free-living animals. Many wild bees are solitary, meaning they do not live in colonies.

These life history differences between wild and managed pollinating insects are important to understand, as their specific life cycles and habitats face different threats. Wild insect pollinators are declining in part due to urbanization and habitat fragmentation (Cunningham, 2000). Previous research has indicated urbanization is associated with reduced ecosystem function (Eigenbrod et al. 2011, Thompson et al 2017). Wild insects are also sensitive to changes in weather and climate, which can impact them directly - as well as indirectly through impacts on their foraging plant hosts (Ogilvie et al. 2017). Honey bees and managed pollinators are often situated in agricultural landscapes for their crop pollination services. In turn, they face threats from disease spread within colonies and the application of pesticides, notably neonicotinoids recently (Vanbergen et al. 2013). Yet wild bees may also be threatened by honey bees (Thomson 2016). As such, the trajectories of managed pollinator populations are not necessarily accurate representations of the health and trajectory of wild insect pollinators (Wood et al. 2020).

Supporting biodiversity of pollinators can better support ecological function. Diverse pollinator assemblages are associated with increased pollination events and efficacy, due to complementary behavior, morphology, and resource use (Bluthgen and Klein 2011, Albrecht et al. 2012). Understanding non-bee pollinating insects has been identified as a key area for research to fully understand pollinating species, especially for non-crop plants (Ollerton et al. 2011, Mayer et al. 2011). For example, beetles (Order Coleoptera) are critical insect pollinators, widely estimated to visit the majority of flowering plants. And while no agricultural crops in the United States are pollinated by beetles, dozens of native plant species are (Kevan and Baker 1983).

In recent years, scientists have encouraged policymakers to address threats to insect pollinators using legislation and other policy tools. This is in large part due to the recognized importance of pollination for crop plants. For example, native insects provided 3 billion dollars worth of crop pollination services between 2001 and 2003 in the US (Losey and Vaughan 2006). Many agricultural plants used by people and livestock as food require insect pollination, an estimated 35% of temperate and tropical food crops according to one estimate (Klein et al. 2011). Additionally, there is increasing evidence that insect populations are declining globally, including pollinating insects (Kluser et al. 2007). For example, bumble bees (*Bombus* spp.) have been documented to be declining across North America (Cameron et al. 2011). While occurrence isn’t necessarily directly correlated with service provision, there is concern that declines in insect pollinator populations could indicate corresponding declines in pollination. Under changing climatic conditions, there could be further declines in pollination, through compounding and synergistic issues, such as phenological mismatches of plants and insects. As such, scientists have written policy targets for pollinator and insect conservation (Harvey et al. 2020, Samways et al. 2020)).

In turn, policies have been created or updated to more explicitly include pollinators or consider impacts on pollinators. For example, an international collaboration coalesced into the Promote Pollinators partnership in 2016, and it now has 30 nations involved (Promote Pollinators n.d). The goal is to advance pollinator initiatives in the member countries. In the European Union, agricultural policy was updated to consider pollinator habitat within agricultural landscapes that align with producer management (Cole et al. 2020). In the US, federal initiatives and legislation that addressed issues related to pollinating insects through agricultural legislation, such as the Food, Conservation and Energy Act of 2008 and Agriculture Act of 2014.

Governments at various administrative levels have also created and passed pollinator protection plans as one of the policy responses. These plans are distinct from embedding considerations of pollinating insects into pesticide or agricultural policies; they center conservation of pollinators. In the United States (US), the federal government created the Pollinator Health Task Force in the executive branch in 2014, via presidential memorandum (The White House 2014). That task force then released the ‘*National strategy to promote the health of honey bees and other pollinators*’ in 2015 (The White House 2015a). It was a collaborative executive branch effort to sync up activities within the federal government to address threats to pollinators with three goals. The goals were to reduce honey bee colony losses to 15% or less annually, increase the monarch butterfly (*Danaus plexippus*) population to 225 million, and to restore 7 million acres of pollinator habitat (The White House 2015a). In practice, it focused heavily on crop pollination and crop pollinators like honey bees. While other wild pollinating insects are mentioned in the federal strategy, the only other focal taxa in the strategy with a goal is the monarch butterfly. Along with the strategy, a “Pollinator Research Action Plan” was developed and released in 2015, to address gaps in knowledge that the task force had identified (The White House 2015b). The plan itself, as a strategy document and not legislation, does not provide funding or have enforcement mechanisms. However, it explains an overarching plan to use existing agencies and structures to achieve a broader mission.

In the US federal government strategy release in 2015, US state governments were encouraged to create their own state-level plans to address pollinating insect conservation. The strategy document itself mentions the federal government developing the initiative in part to lead by example for states and other partners, as well as including coordination with and outreach to non-federal entities (The White House 2015a). Just like the US federal government, many US states already had initiatives and legislation that included attention to insect pollinators.

Our objective was to assess the US state pollinator plans developed and released, to understand how well the plans were designed. Since their stated goal is to promote pollinating insect conservation, we wanted to know whether the state plans are using best practices for evidence-based science policy. The results are critical for understanding if and how pollinator protection plans are contributing to addressing the substantial threats faced by wild and managed insect pollinators. Additionally, the value of this approach is that while this analysis focuses on state level plans, the issues and analytical frameworks scale to all levels of government.

## Methods

### Search strategies

To determine if US states had a state pollinator plan, we searched the phrases “[state name] pollinator plan”, “[state name abbreviation] pollinator plan”, “[state name] AND pollinators site:.gov”, “[state name abbreviation] AND pollinators site:.gov” into the Google search engine in October 2018. If those did not yield results, we went to the state government website and manually searched through the state agricultural agency website. We also tried using search features within state government and state agricultural agency websites. We re-checked the availability of plans in January 2019. When we did find a plan, we downloaded the plan.

### Plan assessment and evaluation

We then performed a close reading and content analysis of all existing US state pollinator plans. This was conducted on all the plans found and downloaded, using a specific list of questions (Appendix 1). We then used three specific frameworks to evaluate and score the information we extracted from the plans. The three frameworks were the United States Environmental Protection Agency (US EPA) managed pollinator protection plan (MP3) guidance (SFIREG 2019), the Pew-MacArthur Results First Initiative evidence-based state policymaking framework (The Pew Charitable Trusts 2017), and the United States Department of Interior (US DOI) adaptive resources management (ARM) framework (Williams et al. 2009). Our analysis assessed each plan using these frameworks to get a multi-dimensional perspective on whether the state pollinator plans are aligned with best practices to potentially achieve their goals of insect pollinator protection and conservation.

### United States Environmental Protection Agency Managed Pollinator Protection Plan (EPA MP3) framework

The US federal plan to protect pollinators specifically included encouragement of states to adopt pollinator plans that matched US EPA guidance for managed pollinators (The White House 2015a). The framework that the US EPA specifically encouraged states to adopt was managed pollinator protection plans (MP3s). The US EPA MP3 framework was designed in collaboration with two stakeholder groups, the Association of American Pesticide Control Officials (AAPCO) and an AAPCO committee called the State FIFRA Issues, Research and Evaluation Group (SFIREG) (EPA OIG 2019). These plans are focused on managed pollinators, which generally refers to honey bees (*Apis mellifera*), one of the two focal species in the federal pollinator plan due to its importance for US agriculture. Very rarely are other pollinating insects kept as managed pollinators; when they are, they can include some bumblebee (*Bombus* spp.) taxa. The contents of MP3s thus focus mainly on issues relevant to managed species in mostly agricultural settings, as opposed to issues facing wild pollinators. According to guidance released by SFIREG, the MP3s for states should include six main elements to match the EPA guidance (SFIREG 2015). These elements are: public stakeholder participation, methods for stakeholders to communicate with each other, use of best management practices (BMPs) to reduce bee exposure to pesticides, an outreach plan, a way to measure the plan’s effectiveness, and a process to review and update the plan. We scored the state pollinator plans to determine their alignment with this framework using these criteria (Table 1). We scored all the available state plans for these elements whether or not they identified as MP3 plans specifically.

**Table 1.** Scoring system for the EPA MP3 evaluation criteria

| **Criteria** | **Components** | **Score** |
| --- | --- | --- |
| Stakeholder involvement | identifying stakeholders | 0.5 |
|  | whether stakeholders were involved in design | 0.25 |
|  | if beekeepers were involved in plan design | 0.0625 |
|  | if growers were involved in plan design | 0.0625 |
|  | if pesticide applications were involved in design | 0.0625 |
|  | if broader public was involved in plan design | 0.0625 |
| Communication Plan | whether communication plan was included | 1 |
| Pesticide Best Management Practices | whether BMPs for reducing pesticide exposure to managed bees was included in plan | 1 |
| Outreach Plan | whether outreach plan was included | 1 |
| Measurement | was there a plan to measure effectiveness | 1 |
| Updates | did plan mention future updates and versions | 0.25 |
|  | was there specific plan to update the document | 0.5 |
|  | was there funding for future updates and work | 0.25 |

Within the 6 overall elements of the US EPA managed pollinator protection plan framework, two of the elements (stakeholder involvement, updates) had sub-elements that were assessed and summed to provide the score for that part of the evaluation criteria.

### Pew-MacAthur Results First Initiative evidence-based state policymaking (PEW) framework

The PEW framework comes from an assessment that the organization did of states engaging in systematic use of analysis and evaluation to guide policy and funding decisionmaking (The Pew Charitable Trusts 2017). The goal of this approach is to use limited resources that states have for programs that are most likely to produce positive results and be cost-effective. Their framework for states using evidence based policymaking included six major elements (The Pew Charitable Trusts 2017). Those are: define levels of evidence, inventory existing programs, cost-benefit analysis assessment, report outcomes, target funds, and require action through state law. Another reason to use this specific metric to assess the state pollinator plans is that the US federal pollinator strategy specifically includes the goal of “*improve targeting of interventions*” and “*review the efficacy of land management actions*” within the plan (The White House 2015a). We scored the state pollinator plans to determine their alignment with this framework using these criteria (Table 2).

**Table 2.** Scoring system for the Pew evaluation criteria

| **Criteria** | **Components** | **Score** |
| --- | --- | --- |
| Assessment | Assessment of overall issue | 0.5 |
|  | State-specific assessment of issue | 0.5 |
| Reviewed previous plans | Whether similar existing plans were reviewed | 1 |
| Cost-Benefit Analysis | Was cost-benefit analysis conducted | 1 |
| Outcomes reporting | Does plan commit to reporting outcomes | 1 |
| Funding allocated | Is funding allocated for the plan's actions | 1 |
| Implementation | Whether plan was a state law | 0.5 |
|  | Identification of which agencies are responsible | 0.5 |

Within the 6 overall elements of the Pew evidence-based state policymaking framework, two of the elements (assessment, implementation) had sub-elements that were assessed and summed to provide the score for that part of the evaluation criteria.

### United States Department of Interior Adaptive Resources Management (DOI ARM) framework

The United States Department of Interior (US DOI) is a federal agency responsible for management of natural and cultural resources. Two of its main responsibilities are habitat management for species conservation and stewardship of land and resources (US DOI n.d.). One of the approaches that US DOI uses to that end is adaptive resources management (ARM). Adaptive resources management is a systematic approach to rigorously, iteratively improve management by learning from outcomes and updating management practices. In their own internal technical guide to applying ARM within the DOI, they identify six main elements of the ARM management process (Williams et al. 2009). These are a cycle that begins with assessing the problem, designing the intervention, implementing the plan, monitoring, evaluating the outcomes, and adjusting the plan based on the evaluation. Another critical reason to include this metric is that the US federal pollinator strategy from 2015 specifically mentions “*engage adaptative management strategies*” within the plan itself (The White House 2015a). We scored the state pollinator plans to determine their alignment with this framework using these criteria (Table 3).

**Table 3.** Scoring system for the US DOI ARM evaluation criteria

| **Criteria** | **Components** | **Score** |
| --- | --- | --- |
| Assessment | Assessment of overall issue | 0.5 |
|  | State-specific assessment of issue | 0.5 |
| Design | were stakeholders involved in design | 0.5 |
|  | were specific goals identified | 0.5 |
| Implementation | identifying implementation goal area | 0.5 |
|  | identifying where implementation is happening | 0.1667 |
|  | identifying who is responsible for implementation | 0.1667 |
|  | specifically identifying habitats for implementation | 0.1667 |
| Monitor | whether plan includes monitoring | 1 |
| Evaluation | does plan evaluate outcomes | 1 |
| Adjust | was there a plan to adjust based on evaluation | 1 |

Within the 6 overall elements of the US Department of the Interior adaptive resources management framework, three of the elements (assessment, design, implementation) had sub-elements that were assessed and summed to provide the score for that part of the evaluation criteria.

## Results

### State pollinator plans

As of early 2019, 31 of 50 states had developed and released a state pollinator protection plan (Figure 1). Of those, about a third (n=12) mentioned the 2014 US presidential memorandum as a motivation for their state pollinator plan, indicating that many states did create their plans as a result of encouragement from the federal initiative. Most of the state pollinator plans were released in 2016 (n=16), shortly after the US federal pollinator plan was released. Two states (Maine and Wyoming) released their plan in 2015, the same year as the federal plan, and one state (North Dakota) released their plan in 2014, the same year as the federal pollinator task force was established. The rest were released in 2017 (n=8) and 2018 (n=4). Regardless of the release date or start year of the plans, most did not identify an end date. Only two of the plans included a specific end year; Delaware and Missouri’s plans said it would expire in 2019. Massachusetts’ plan said it was intended for a 5-10 year timeframe. Five states said they would revisit the plan annually. Most of the plans (n=25) did not include information on if and how to update the plan, yet almost the same number (n=23) mention “update” in the plan itself.

**Figure 1.**
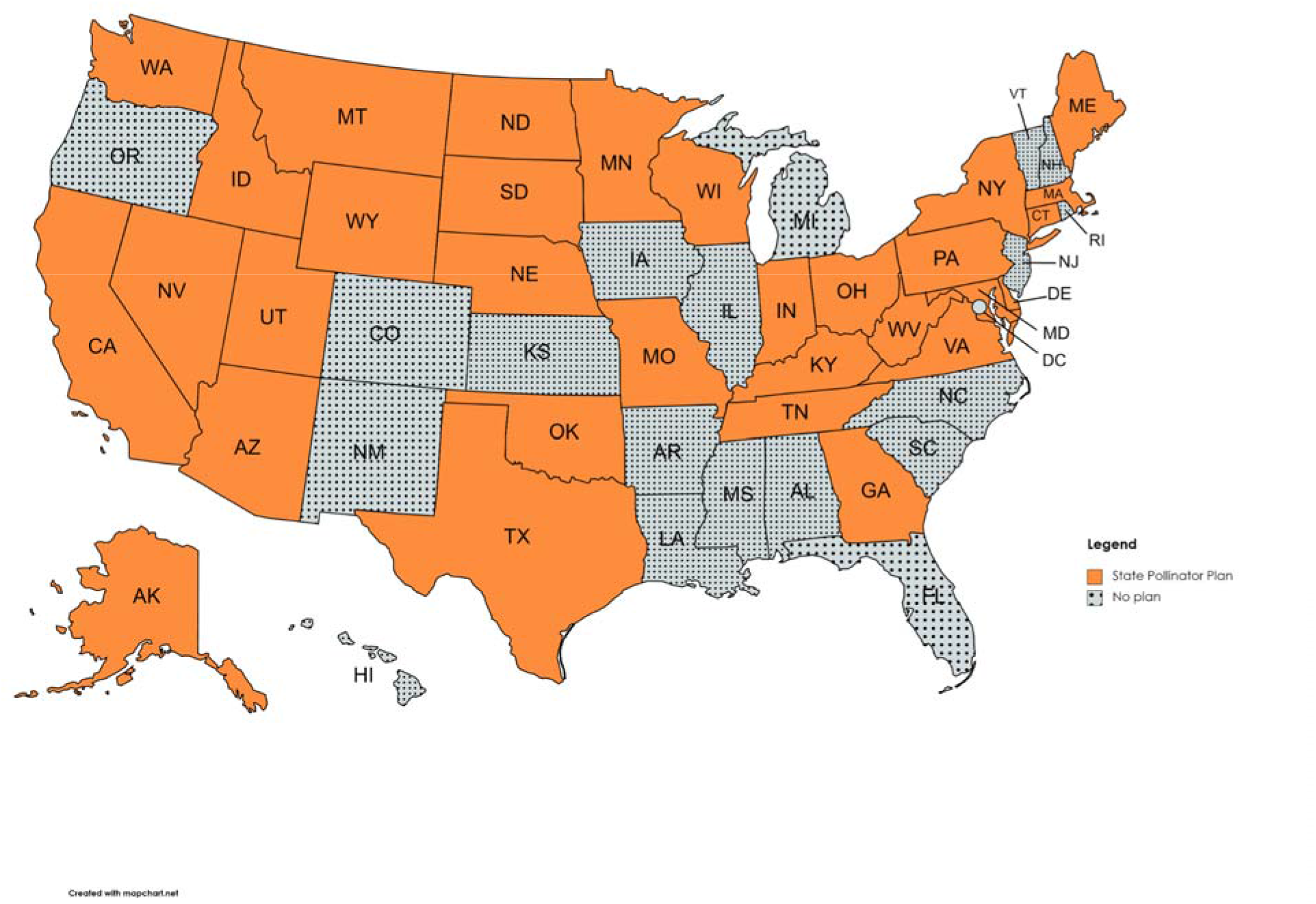
Map of US states with pollinator plans. A map of the United States states indicating which states had a state pollinator plan publicly released during the study period. Orange coloration indicates states that have a plan and grey coloration with dots indicates states that do not have a plan.

Most of the plans were written by the state department of agriculture (n=14) or in collaboration with the department of agriculture (n=5). This reflected a close but not perfect alignment with which plans were managed pollinator protection plans (MP3s). Of the plans, just over a third (n=13) are MP3s (Figure 2). This is an indication they are focused on honey bee pollinator protection issues, one of the two focal taxa of the federal plan. Monarch butterflies (*Danaus plexippus*) are the other focal taxa in the federal plan, and they were specifically included in less than half of the state pollinator plans (n=14).

**Figure 2.**
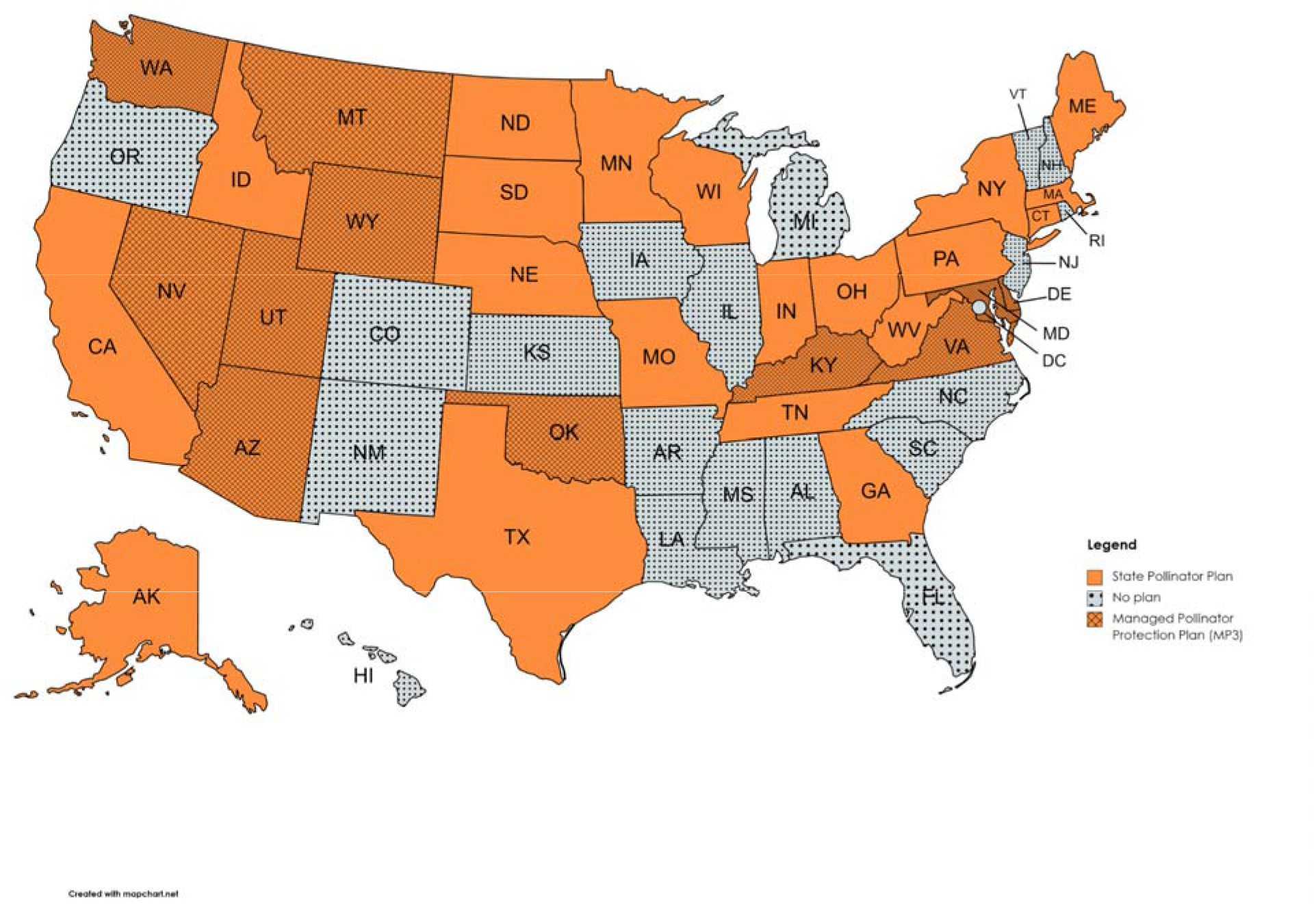
Map of US states with MP3s versus other types of pollinator plan. A map of the United States states indicating which states had a state pollinator plan specific to managed pollinators versus a broader state pollinator plan during the study period. Orange with hatch marks indicates a managed pollinator protection plan (MP3). Orange with no hatch marks indicates a broader state pollinator plan. Grey with dots indicates a state with no plan during the study period.

### Alignment with EPA MP3 guidance

No state plan was completely aligned with the EPA MP3 guidelines (Figure 3, Figure 4). The states that scored highest (above 75% of possible points per Table 1) were Nevada, Wisconsin, Massachusetts, Nebraska, New York, Indiana, Delaware, and California. 20 states scored half or more of the possible assessment points under the MP3 guidance. The alignment of which states scored the highest in matching the EPA MP3 elements was not strongly associated with whether the state had a plan labeled as a managed pollinator protection plan (Figure 2).

**Figure 3.**
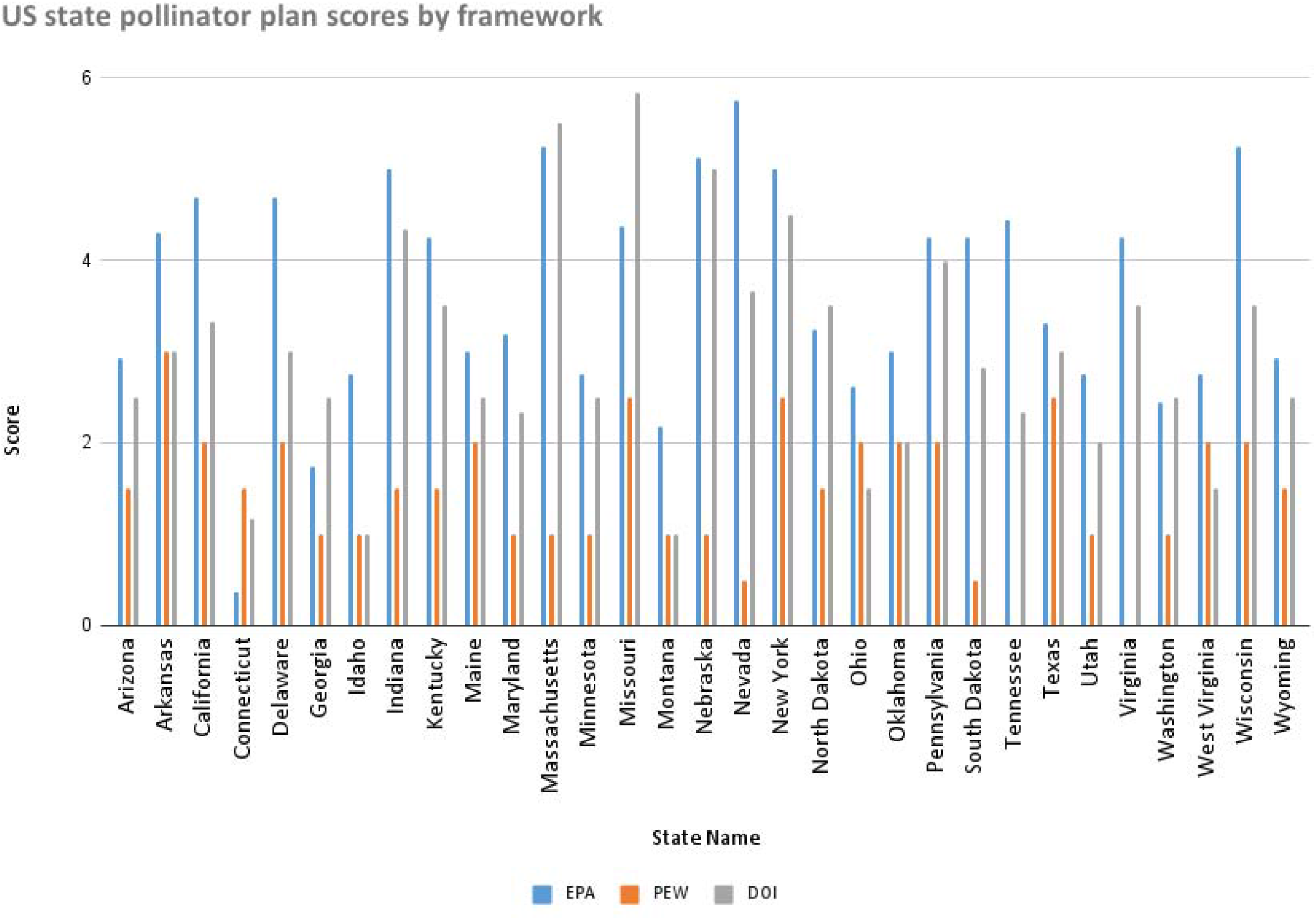
US state pollinator plan scores by framework. Bar chart of the United States states and how they score on the 3 evaluation metrics on a scale of 0 (lowest) to 6 (highest). The blue bar represents the EPA MP3 score, the orange bar represents the PEW framework score, and the grey bar represents the DOI ARM score.

**Figure 4.**
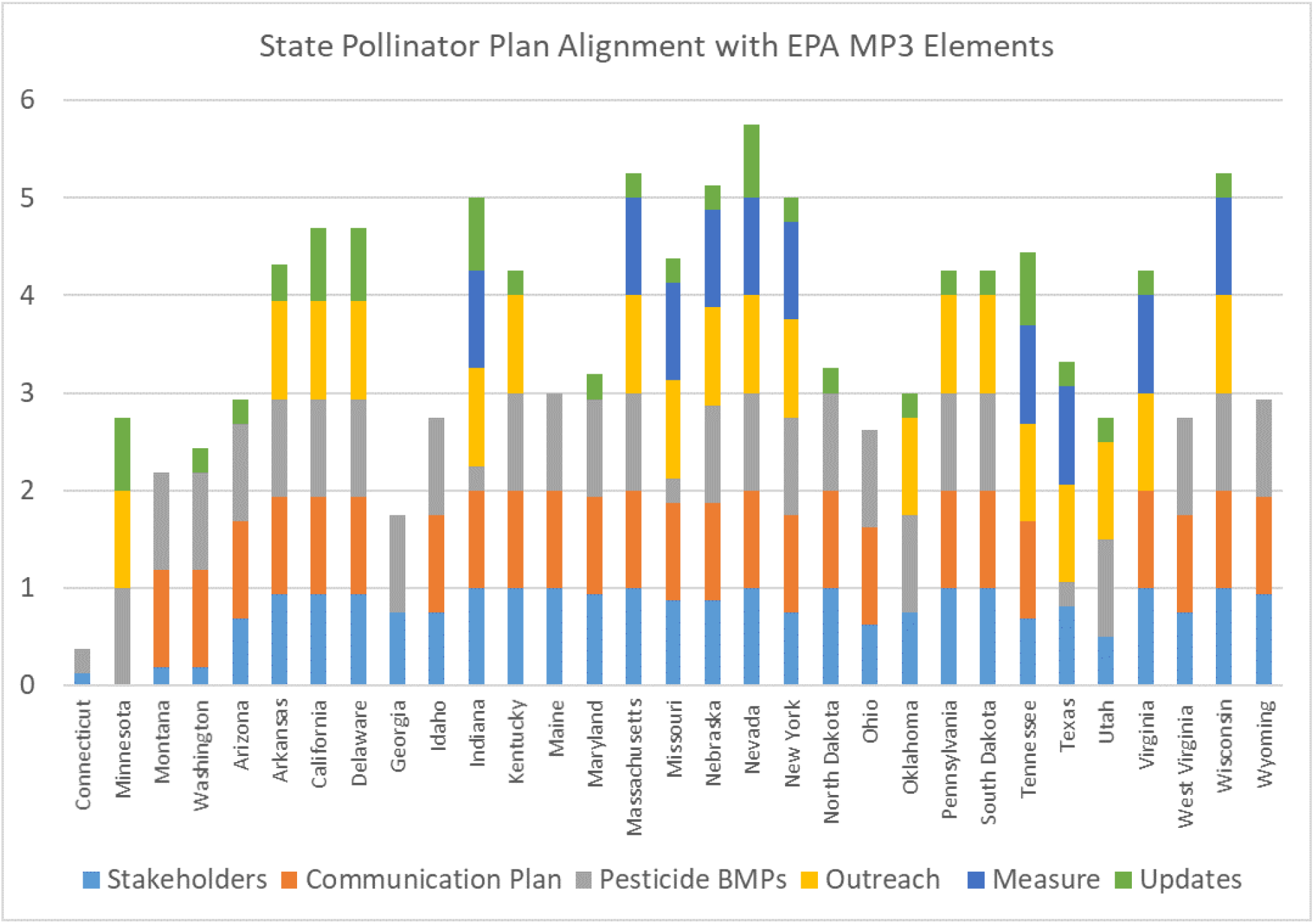
State Pollinator Plan Alignment with EPA MP3 Elements. This figure shows the breakdown of alignment with the EPA MP3 guidance by element for all state pollinator plans. All states were scored, even if the plan was not labeled explicitly as an MP3. State figures appear in alphabetical order; states without plans are not included. The axes match the scoring (Table 1) and appear bottom to the top as how plans scored on: stakeholder involvement, communication plan, pesticide best management practices (BMPs), outreach plan, measurement, and updates.

The EPA MP3 guidance specifically wanted to see beekeepers, growers, pesticide applicants, and the broader included as stakeholders. The overwhelming majority of state plans, whether or not they identified as MP3 plans, discussed all of these stakeholder groups explicitly. The majority of plans (n=27) identified relevant stakeholders in the plan. Yet fewer (n=17) discussed these stakeholders being involved in the design of the plan. Another key element of the MP3 guidance was including a communication plan for stakeholders. The vast majority of states with pollinator plans (n=25) specifically discussed the communication plan. Fewer state plans (n=19) included specific outreach plans about the pollinator protection plan.

A unique dimension of the EPA MP3 guidance compared to the other evaluation frameworks in our analysis was the requirement to include best management practices (BMPs) for reducing managed pollinator exposure to pesticides. Only two plans did not do this all. Yet only a third (n=11) plans included discussion of measuring the impacts of the plan.

All states, even ones explicitly releasing plans billed as MP3s, scored poorly on the update related evaluation criteria. While many plans mentioned needed to update the plan in the future (n=23), few had specific timelines or other information on when or how that would happen (n=6). Additionally, only one state (Arkansas) mentioned funding for a plan update. Even that one state did not mention a specific amount of funding though.

### Alignment with Pew evidence-based state policymaking framework

No state pollinator plans scored highly when evaluated with the Pew evidence-based state policymaking framework (Figure 3, 5). No state received greater than half the assessment points (higher than 3 out of 6). Only one state (Arkansas) received exactly half the possible evaluation points by including information that matched half the required elements under the PEW framework. Every other state (n=30) scored 2.5 out of 6 or lower; two state plans (Tennessee, Virginia) scored a zero using this assessment tool.

**Figure 5.**
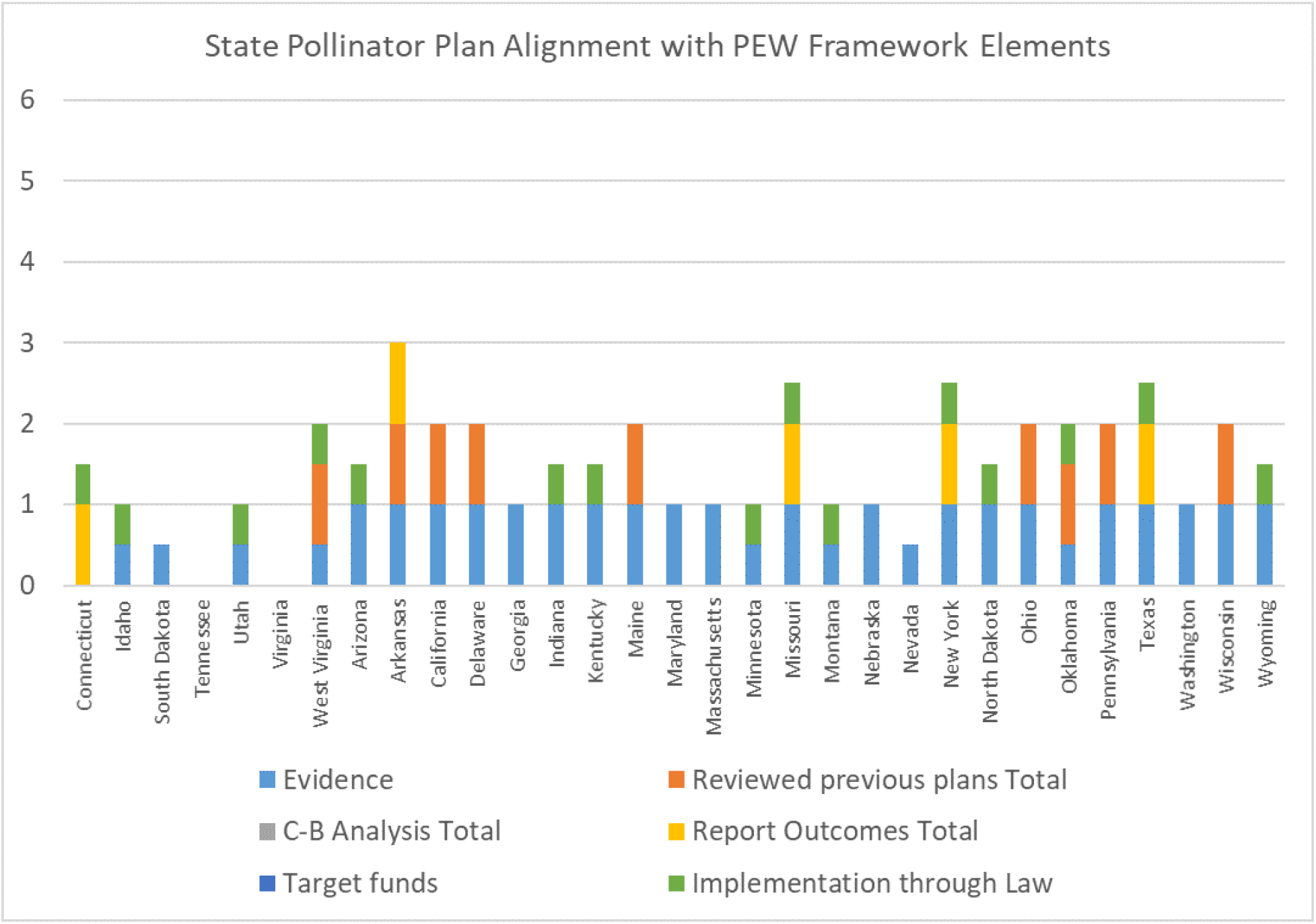
State Pollinator Plan Alignment with PEW Elements. This figure shows the breakdown of alignment with the PEW framework by element for all state pollinator plans. State figures appear in alphabetical order; states without plans are not included. The axes match the scoring (Table 2) and appear from the bottom to the top as how plans scored on: assessment, reviewed previous plans, cost-benefit analysis, outcomes reporting, funding allocated, and implementation as state law.

The PEW framework for evidence-based state policymaking requires assessment of the problem to be addressed by policy and reviewing existing, related programs. Most state plans (n=24) had assessed the baseline situation and the same number specifically included state-level information about the status of pollinating insects the plan was to address. Most states (n=29) included some review of what levels of evidence meant in the plan. Yet many states did not review their own existing state initiatives or policies, whether legislation or other policy tools, in the plan. Only 9 states mentioned that information in their pollinator protection plan. In turn, a unique element of the PEW framework is the requirement to include a cost-benefit analysis as part of the plan design. No state included this in their plan as part of plan design discussion.

Monitoring and evaluation is another key part of the PEW framework. Yet most state plans also scored poorly there. Only 5 state plans included mention of reporting outcomes of the plan. No plans included funding to achieve the goals or implement the activities mentioned in the plan. Yet almost half of the plans (n=14) did identify who was supposed to implement the plan. Only one state (Connecticut) enacted their state pollinator plan as a state law.

### Alignment with DOI ARM framework

Many state pollinator protection plans did not align well with the DOI ARM framework (Figure 3, Figure 6). While one state (Missouri) had a near perfect score in terms of including all the ARM elements, most state plans (n=28) scored less than 75% of the possible points (4.5/6) using this assessment tool. Massachusetts and Nebraska also scored very highly. Yet 15 state plans scored less than half the possible points (3/6) under this framework.

**Figure 6.**
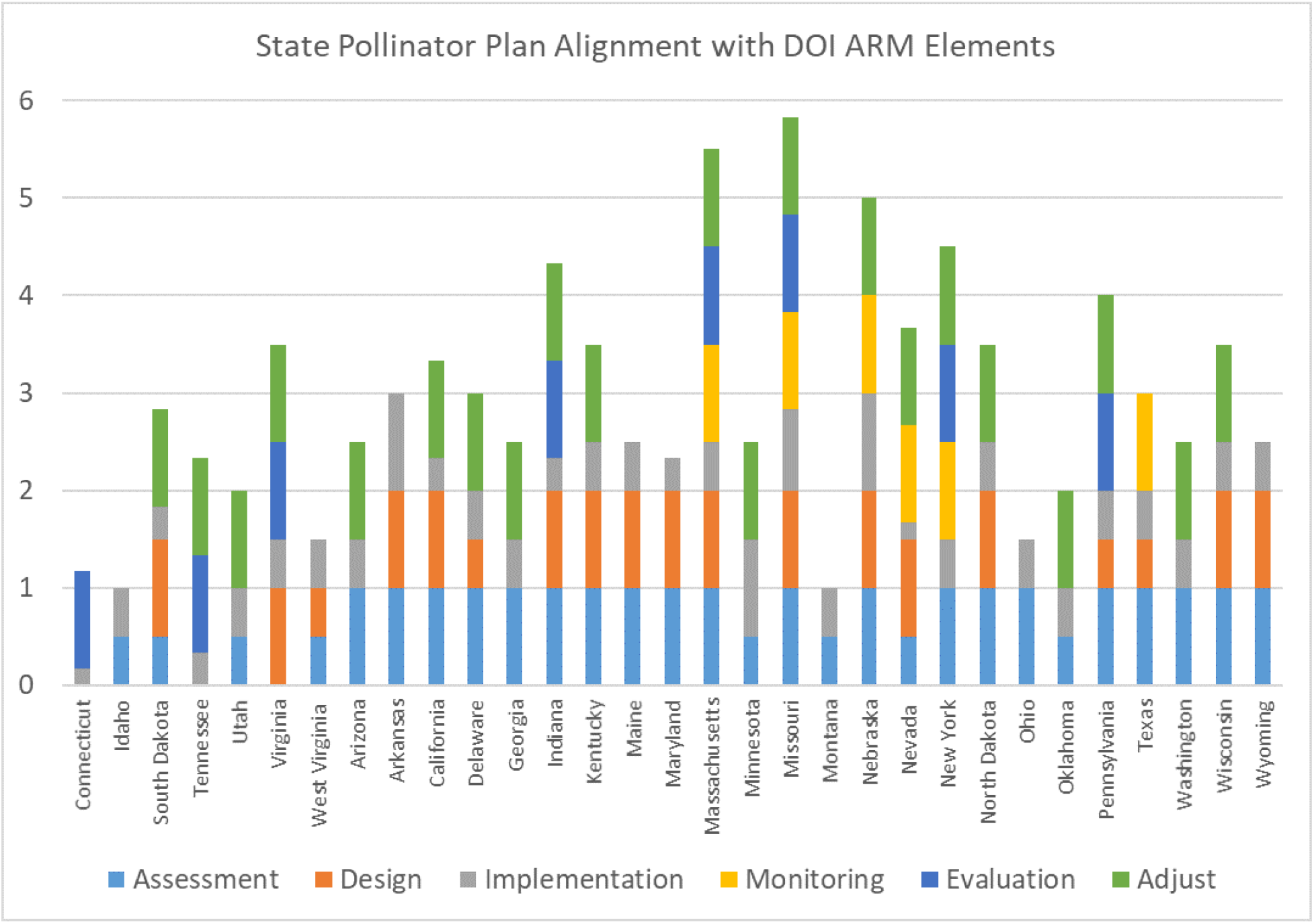
State Pollinator Plan Alignment with DOI ARM Elements. This figure shows the breakdown of alignment with DOI ARM by element for all state pollinator plans. State figures appear in alphabetical order; states without plans are not included. The axes match the scoring (Table 3) and appear from the bottom to the top as how plans scored on: assessment, design, implementation, monitoring, evaluation, and adjustment.

The disconnect between state plans and ARM elements was mostly seen in the lack of monitoring, evaluation, and adjustment in state plans. Very few states (n=6) mentioned monitoring as part of their pollinator plan. And while 8 states included mention of evaluating the plan outcomes, the majority of state plans said they would adjust the plan in the future based on results (n=21).

Yet as seen in the PEW framework results, most states with plans did have plans that assessed conditions before the plan, including at the state-level. And most plans included stakeholders in their design process (n=18) and had design process elements that aligned with ARM principles (n=16). Of all of the state plans, 71% (n=22) also included specific, defined goals to achieve with the plan.

Results were more mixed with specifics in the plan about implementation details. Only 3 state plans (Minnesota, Nebraska, Arkansas) included all of the implementation information (goal area, where, who, and which habitats). Most plans (n=27) did not include implementation goal areas. Every plan included information on who was responsible for implementation and almost all included mention of which habitats implementation was slated for (n=28).

## Discussion

Overall, we found that US states are very varied in whether or not they have state pollinator protection plans and what’s in the plans. Additionally, how well those plans align with best practices for evidence based policymaking, using three distinct frameworks, also varied significantly. Of the 31 states with a state pollinator plan, Connecticut was the state with the lowest total score. The state with the highest overall scores, across the three frameworks, was Missouri. Most states did not score highly consistently across the frameworks.

While previous work examined pollinator-relevant legislation passed by states and found over a hundred in the past two decades (Hall and Steiner 2019), we found that only one state pollinator plan was passed as legislation (Connecticut). This means the rest of the states did not require action through state law, which the PEW framework considers a benchmark of effective state policy. Thus this analysis is a critical contribution to the knowledge on this topic, as assessment of pollinator policies that are laws leaves out the other 30 state pollinator plans. Yet there is significant overlap of the states that have both a state pollinator plan and have legislation that addresses pollinators (Hall and Steiner 2019. The disconnect between plans and legislation has material consequences. While we found that no state plans included funding in the plan itself, many pieces of passed legislation about pollinators in the same states did include funding allocations, for research or implementation (Hall and Steiner).

This analysis also provides a critical, independent assessment of the state pollinator plans. According to the US EPA, as of January 2018, 45 states “had developed or were developing” state-level MP3 plans (US EPA OIG 2019). Then in 2019, AAPCO surveyed representatives from each state about the contents of their MP3 plan. The self-reporting from states on alignment with the MP3 guidance is much higher than the results we found in our analysis (AAPCO 2019). One area where the AAPCO survey and our results did agree was the reporting on updating and reviewing plans. Respondents mostly reported (~75%) that they were not meeting to review and update the MP3 plans (AAPCO 2019). Additionally, despite the involvement of EPA in designing the MP3 elements and in co-designing an evaluation survey of those plans in 2019, the EPA itself, via an EPA Office of Inspector General 2019 audit report, concluded that it did not have a strategy to evaluate the impact of the state MP3 plans (US EPA OIG 2019). This finding matches the plans themselves, which mostly did not include specific monitoring and evaluation plans - nor funding to implement the plan or update it in the future.

Pollinator conservation under global urbanization and a changing climate is full of uncertainty. Including iterative monitoring and management, such as ARM, is designed specifically to deal with the uncertainty of science at landscape scales and management innovation. Yet we found most plans lacked specifics and resources for planning beyond the release of the plan itself. This finding is not hopeful for the goal of achieving the intended plan outcomes - or being able to document them if achieved. While ARM is not the only management framework that incorporates uncertainty, policy about science must address complexity and uncertainty (Renn et al. 2019).

While this analysis has revealed major patterns in US state pollinator plans and limitations in the plans, this approach has limitations. It is possible that our assessment is conservative and underscores the state plans. This could happen if the state plans included elements in the evaluation frameworks but did not discuss them in the plan itself. This seems highly possible for elements released to stakeholder involvement and design. There are also many other potential ways to evaluate the pollinator protection plans. For example, a team of scientists released ten policy items that would promote pollinating insect conservation (Dicks et al. 2016). It is possible that the state pollinator plans would score higher or differently if assessed using those criteria instead. Yet an analysis of US legislation related to pollinators found most laws lacked those elements (Hall and Steiner 2019). Additionally, states may have now released updated plans that address some of the limitations discussed herein during the analysis. The revised plans may score higher. For example, New York released a “plan update” in 2020 that was not considered in this analysis (NYS 2020). Moreover, two states had written into their plan their intention to revise after 2019. Future research on these first sets of plans and subsequent updates could reveal if and how plans evolve.

### Conclusions

Many US states designed and implemented state-level pollinator protection plans after the US federal government released a pollinator strategy for national agencies. Yet the state pollinator plans do not necessarily contain the elements and resources that are thought to best position the states to achieve their intended goals of improved pollinating insect conservation. Many plans did not include funding for implementation, specific evaluation metrics, or concrete plans to revise and update plans based on monitoring and evaluation. These missing elements may impact states’ ability to achieve their conservation goals. Our results indicate that states can improve their pollinator conservation policies to better match evidence-based science policy guidance, regardless of which framework is used.

## Acknowledgments

Funding for this project came in part from a Rochester Institute of Technology College of Science 2018 faculty development grant.

## Data Availability

All the data for this study and analysis is freely available here:

Kaitlin Stack Whitney, & Briana Burt Stringer. (2021). Evaluation of US State Pollinator Plans Using Three Evidence-Based Policymaking Frameworks [Data set]. Zenodo.

http://doi.org/10.5281/zenodo.4917595.

## Appendices

## Appendix I. Full list of data collected from the state pollinator plans

- If the plan was found (and if so, it was downloaded and saved)
- The year it was written and provided to the public
- The agency that wrote it
- If the state department of agriculture wrote the document, or if they along with another organization wrote it
- Does the plan specify an end date for the plan?
- Does the plan specify a timeframe within the plan to address when tasks/monitoring will be completed?
- Does the plan present quantitative information on the nationwide issue of pollinator decline/importance?
- Does the plan present quantitative information about the state specific issue of pollinator decline/importance?
- Does the plan describe how the plan was first designed/crafted?
- Does the plan mention that the organizers sought out groups to help determine what needs to be included?
- Does the plan mention specific goals and/or objectives that are achievable?
- Does the plan have specific goals/objectives on how the plan will be implemented within the state? This could include creating jobs, funding for projects, specifying acreage, allocating resources for a specific task, etc.
- Whose land is the plan supposed to be implemented on (e.g. state agencies, private landowners)? - Could be one, multiple, or none.
- Does the plan specify a specific amount of land (acreage) that they hope to implement a part of the plan on? State specific acreage if mentioned.
- Does the plan specify who will do the work of implementing the plan? If so, which agencies or organizations/people
- Does the plan include which specific habitats implementation of the plan will occur (e.g. cities, parks, forests, agriculture, lawns, gardens, fields (meaning agricultural, typically crop)? Specify all.
- Does the plan include a plan/goals to monitor the progress of the plan, as it is being implemented?
- Does the plan have a plan to review the outcome of the project to determine if the management plan was successful?
- Does the plan state an evaluation metric?
- Does the plan have a plan to adjust/change/update the plan as they monitor and evaluate the outcomes?
- Does the plan mention stakeholder participation as part of the plan?
- Does the plan take into account stakeholder review while designing/writing the plan?
- Does the plan mention beekeepers as a stakeholder?
- Does the plan mention agricultural “growers” as a stakeholder?
- Does the planmention “landowners” (versus agricultural workers) as a stakeholder?
- Does the plan mention the public as a stakeholder?
- Is communication between stakeholders a goal of the plan?
- What method of communication between stakeholders do they include as a goal of the plan? (e.g. meetings, discussions, etc.)
- Does the plan include the best management practices (BMPs) for reducing pesticide exposure?
- Does the plan specify a plan for outreach?
- Are managed pollinators mentioned in the plan? Mentioned (M),
- Is the plan a MP3 (managed pollinator protection plan) specifically?
- Does the plan plan on monitoring/measuring specifics to determine if the MP3/plan is effective and will accomplish its goals?
- Does the plan mention wanting to update the plan?
- Does the plan actually have a plan to periodically review and update the plan?
- Does the plan mention a year that they will evaluate the plan for updates? If so, what year?
- Is there specific funding allocated for plan updates?
- How much funding in US dollars is that amount?
- Are levels of evidence and success defined in the plan?
- Does the plan specify if they reviewed previous or similar plans before drafting their plan?
- Does the plan use a cost-benefit analysis - to compare cost of program/plan compared to the economic returns?
- Will they report the outcomes of the plan? Specifically does the reporting go beyond what is already required by other state laws/regulations (such as reporting pesticide incidents or managed hive locations etc.)
- Does the plan include targeted funds to achieve the plan?
- Is the plan a law?

